# Biomechanics Characterization of Autonomic and Somatic Nerves by High Dynamic Closed-Loop MEMS force sensing

**DOI:** 10.1101/2023.04.13.536752

**Authors:** María Alejandra González-González, Hammed Alemansour, Mohammad Maroufi, Mustafa Bulut Coskun, David Lloyd, S. O. Reza Moheimani, Mario I. Romero-Ortega

## Abstract

The biomechanics of peripheral nerves are determined by the blood-nerve barrier (BNB), together with the epineural barrier, extracellular matrix, and axonal composition, which maintain structural and functional stability. These elements are often ignored in the fabrication of penetrating devices, and the implant process is traumatic due to the mechanical distress, compromising the function of neuroprosthesis for sensory-motor restoration in amputees. Miniaturization of penetrating interfaces offers the unique opportunity of decoding individual nerve fibers associated to specific functions, however, a main issue for their implant is the lack of high-precision standardization of insertion forces. Current automatized electromechanical force sensors are available; however, their sensitivity and range amplitude are limited (i.e. mN), and have been tested only *in-vitro*. We previously developed a high-precision bi-directional micro-electromechanical force sensor, with a closed-loop mechanism (MEMS-CLFS), that while measuring with high-precision (−211.7μN to 211.5μN with a resolution of 4.74nN), can be used in alive animal. Our technology has an on-chip electrothermal displacement sensor with a shuttle beam displacement amplification mechanism, for large range and high-frequency resolution (dynamic range of 92.9 dB), which eliminates the adverse effect of flexural nonlinearity measurements, observed with other systems, and reduces the mechanical impact on delicate biological tissue. In this work, we use the MEMS-CLFS for *in-vivo* bidirectional measurement of biomechanics in somatic and autonomic nerves. Furthermore we define the mechanical implications of irrigation and collagen VI in the BNB, which is different for both autonomic and somatic nerves (∼ 8.5-8.6 fold density of collagen VI and vasculature CD31+ in the VN vs ScN). This study allowed us to create a mathematical approach to predict insertion forces. Our data highlights the necessity of nerve-customization forces to prevent injury when implanting interfaces, and describes a high precision MEMS technology and mathematical model for their measurements.

## Introduction

Micro-electromechanical systems (MEMS) are miniaturized devices with dimensions ranging from less than one micrometer to several millimeters, featuring both mechanical and electrical components ^1–3^. MEMS technology has offered innovative devices in various domains of science and technology. The use of MEMS in biological systems ranges from measuring force-displacement characteristics of soft tissues (i.e. liver and maxillofacial muscle) during laparoscopic procedures ^4^ to the investigation of mechanical properties of soft and hard tissue (e.g. muscle and bone) for surgery training ^5^. Existing MEMS devices are mostly unidirectional ^6, 7^, which limits their application to biologically samples (e.g. neuronal tissue) due to their highly dynamic nature, the difficulty of access, limited measurement range and resolution, as well as limited bandwidth. Consequently, mechanical parameters obtained from neural tissues have been measured *in-vitro* from explanted or frozen/thawed tissues ^8–10^, which modifies the biomechanical properties of the sample. This highlights the impact of *in-vivo* measurement which in part requires the MEMS force sensor to be able to measure bi-directional forces with a high bandwidth in a closed-loop system on those dynamic targets such as *in-vivo* peripheral nerves.

The study of mechanical properties of peripheral nerve tissue has relevance in microsurgery to prevent tissue damage by reducing collision force on the surgical instrumentation, and in the development of chronic-implantable neural interfaces, such are regenerative prosthesis and intra/extraneural electrodes for electrical stimulation in bioelectronic medicine (i.e. vagus nerve stimulation for the treatment of epilepsy, morbid obesity and intractable depression, or the sciatic nerve to modulate the blood pressure in hypertensive subjects) ^11–14^. The biomechanics of the nerves are dictated by the blood-nerve barrier (BNB), together with the epineural barrier, extracellular matrix, and axonal composition^15, 16^. Limited is the knowledge about the biomechanics of the BNB, and a comprehensive study about the differences between somatic and autonomic nerves is still needed, since the fabrication of neural interfaces requires the use of materials that mimic those mechanical properties, such as stiffness, viscoelastic modulus, and linear elasticity ^17–19^. This will reduce the mechanical mismatch between the neural interface and the tissue, reducing the long-term damage due to inflammation and the growth of fibrotic tissue within the nerve-interface space. Recently we described the design and characterization of a bi-directional MEMS-based closed-loop force sensor (MEMS-CLFS) with a high force sensing range resolution (−211.7 μN to 211.5 μN), and dynamic range (92.9 dB) ^20^. The MEMS-CLFS technology consists of an on-chip electrothermal displacement sensor and electrostatic actuators. The embedded electrothermal sensor achieved a sub-nanometer displacement resolution (0.645 nm) with the assistance of an incorporated on-chip leverage mechanism that amplifies the shuttle beam’s displacement. The force sensor is being used in a closed-loop configuration, while the control loop maintains the null position of the shuttle beam (end-effector) in the presence of the external force. Using the control scheme as well as the high-resolution displacement sensing, the device demonstrated a resolution of 4.74 nN, with a measurement bandwidth of 92.9 dB. Functioning in closed-loop mitigates the non-linearities which could be introduced by flexural mechanism.

In this work we report the use of our bi-directional MEMS-CLFS to measure *in-vivo* the biomechanical properties in a microscale range of both somatic and autonomic nerves of rat (i.e. sciatic and vagus nerve, respectively), with high precision and resolution. We evaluated the blood nerve barrier density distribution in both nerves, and evaluated the correlation with the biomechanics in areas with different degree of irrigation. This study allowed to generate a mathematical model that allows to predict the force required to differentially penetrate autonomic and somatic nerves. This data will benefit the design of neural interfaces and prosthesis to predict the collision forces during microsurgery with the goal of reducing tissue damage.

## Results

### MEMS-CLFS fabrication

The MEMS-CLFS was fabricated using a silicon-on-isolator multi-user MEMS process (SOI-MUMPs, MEMSCAP Inc). The three-mask process starts with a silicon-on-isolator wafer comprising a 25-μm-thick Si device layer, a 2-μm-thick buried-oxide, and a 400-μm-thick handle layer. The microfabricated devices were wire-bonded to a custom-designed printed circuit board (PCB). Figure 1 shows the MEMS-CLFS schematic design (a) and the representation of the external forces inducing the displacement in the internal mechanism of the probe, measured by an electrothermal sensor (Figure a’). The MEMS-CLFS was designed for *in-vivo* penetration of the sciatic and vagus nerve of the rat, with an insertion tip length of 45.5 μm, a total length of 482 μm, and the surface area of the tip of 2x10 μm to be aligned with the longitudinal arrangement of the axons in the nerves during insertion (Figure 1c). The MEMS-CLFS comprises a shuttle beam with a probe at its extremity actuated by two sets of electrostatic actuators that are used for the closed-loop operation of the device. In ^20^ we detailed the characterization procedure of these electrostatic actuators. The induced displacement on the shuttle beam is amplified by a leverage mechanism with an amplification ratio of 2.5 before it is measured by the embedded electrothermal displacement sensor. The structure of this electrothermal displacement sensor is detailed in ^20^. By amplifying the shuttle beam’s motion, the displacement measurement resolution is enhanced. The 1σ-resolution of the electrothermal sensor is reported as 0.645 nm in ^20^. The device has a trapezoidal-shaped parallel plate capacitive structure that can be used to adjust the stiffness of the device. The force sensor is wire-bonded on a custom-designed printed circuit board (PCB) which is connected to a separate signal conditioning PCB using a flexible flat cable. The signal conditioning PCB comprises the necessary circuitry for the actuation signal as well electrothermal sensor readout.

**Figure 1.**
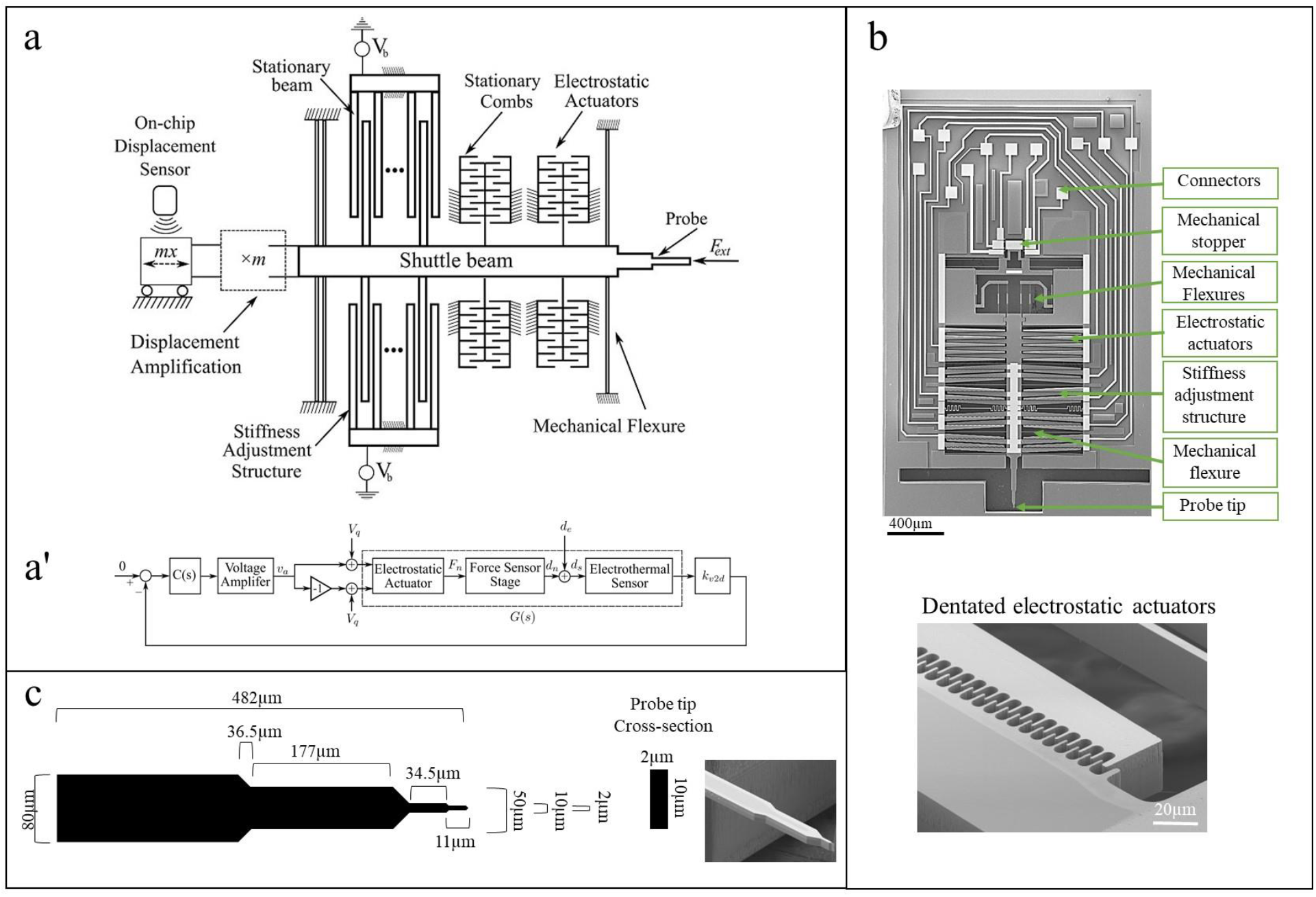
High-dynamic range MEMS-CLFS design. a) Mechanistic design of the MEMS-CLFS. a’) Closed-loop block diagram, the external force induces displacement *d_e_*, on the force sensor stage which is measured by the electrothermal sensor. This external displacement is nullified by the embedded electrostatic actuator that pushes back the stage by *d_n_*. The controller C(s) adjusts the input voltage of the electrostatic actuator v_e_, to the amount that is needed for compensating the external displacement. b) SEM pictures of the MEMS-CLFS final design. c) Schematic representation of the insertion tip with dimensions, and SEM image of the tip probe.

### MEMS-CLFS characterization

The MEMS displacement sensor measures the displacement of the shuttle beam when an external force is applied to the force sensor probe, i.e. the force to be measured when the probe is inserted. Force sensors are conventionally operated in an open loop configuration, where the induced displacement of the shuttle beam is measured and directly converted to force by Hook’s law. However, this approach suffers from flexural mechanisms and nonlinearities. We designed our MEMS-CLFS to be operated in closed-loop. In this configuration, the initial displacement of the shuttle beam is compensated by an actuation force and the probe remains at zero-displacement. The feedback controller sends a command signal to the embedded actuators in order to nullify the external force. This pushes back the MEMS force sensor shuttle beam to its zero-displacement position. In the steady state, the actuation signal *u* is proportional to the external force (Figure 1a-a’). The MEMS force sensor actuators were characterized to convert the actuation signal (in volts) to force (in µN).

### In-plane Static Response

In-plane displacement of the shuttle beam along the x-axis was measured by a laser doppler vibrometer (Polytec MSA-100-3D) and plotted as a function of the actuation voltage (Figure 2a), and the output of the electrothermal sensor was simultaneously recorded. These two parameters were used to calculate the calibration factor of the MEMS-CLFS, the electrothermal sensor output was plotted versus the shuttle beam displacement, and the slope of the resulting curve represents the calibration factor of the device, which is obtained as µm/V (Figure 2b).

**Figure 2.**
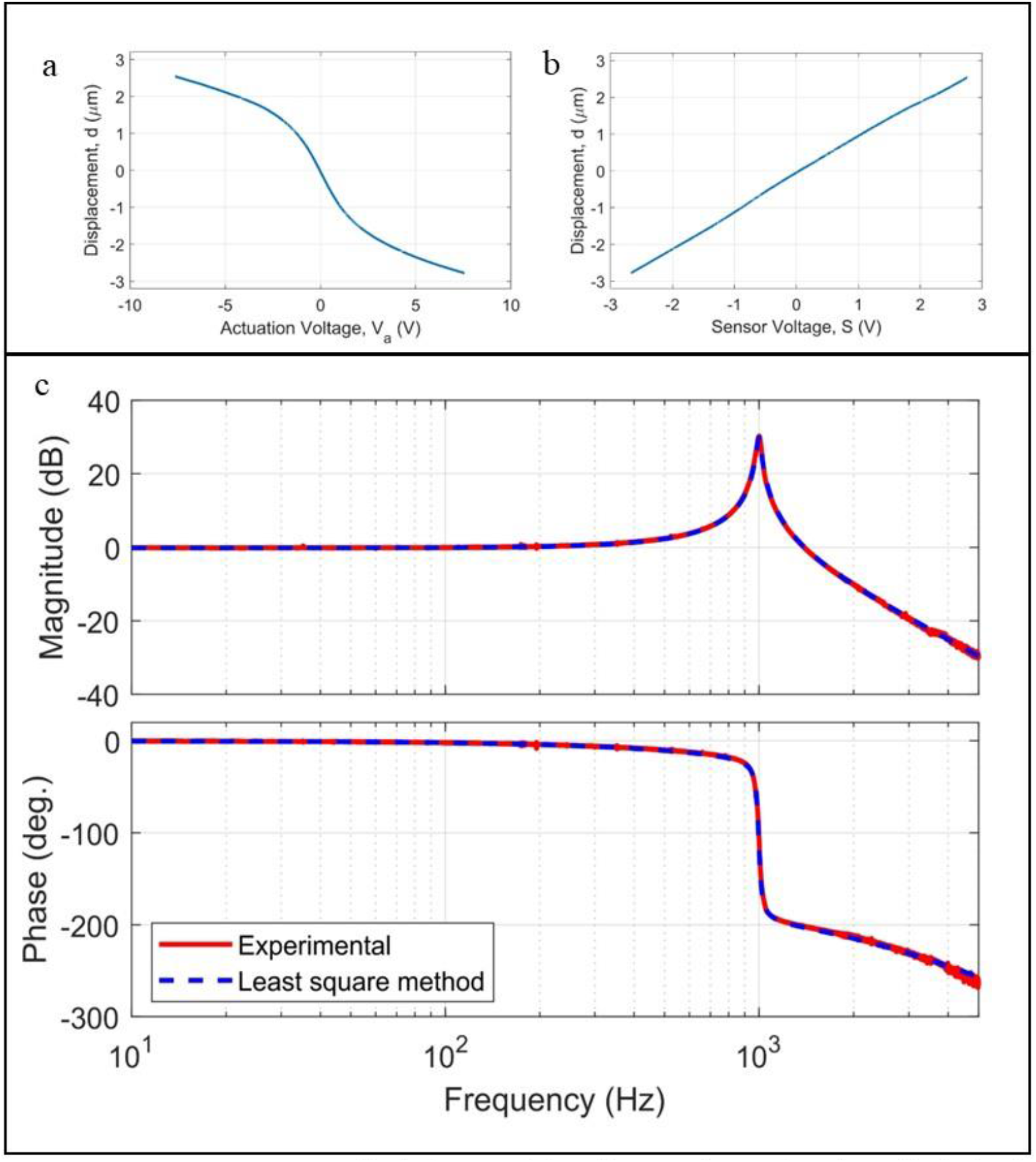
MEMS-CLFS calibration. (a) Displacement of the probe versus the actuation voltage. (b) The shuttle beam displacement as a function of the electrothermal sensor voltage output provides the slope that represents the calibration factor. (c) Experimental frequency response of the force sensor from the actuation voltage input to the electrothermal sensor output along with the identified transfer function response. The frequency response was ‘measured from 10 Hz to 5 kHz.

### Frequency Response Measurement

The frequency response of the MEMS-CLFS was obtained from the actuation input to the electrothermal displacement sensor output, using a chirp excitation signal provided by a CF-9400 ONO SOKKI FFT analyzer. The resonance frequency of the device was obtained as 998 Hz (Figure 2c).

### Calibration and parameter identification

To calibrate the sensor, the MEMS-CLFS actuators were characterized so that they convert the actuation signal (in volts) to the generated force (in N). We previously reported ^20^ that the net force exerted on the shuttle beam by the actuators is:

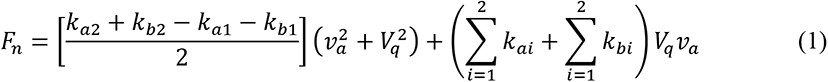

where *ka1,2* and *kb1,2* are actuation coefficients that are defined based on the rate of change of comb capacitances as:

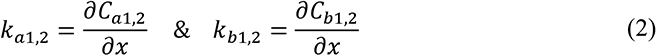

We elaborated on the characterization process in ^21^. An experimental procedure was proposed there with which we can find the four unknown actuation coefficients (i.e., *k_a,_*_1,2_, and *k_b,_*_1,2_) as well as the mechanical stiffness of the force sensor (*k_FS_*). By following the procedure in ^20^, the relation between the actuation voltage and force is experimentally obtained as:

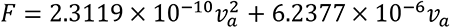

### Control design and implementation

The experimental frequency response (Figure 2c) was used to model the force sensor. A transfer function with one zero and three poles was adopted for this purpose, the coefficients of which were obtained by the least square method. The identified transfer function is represented as:

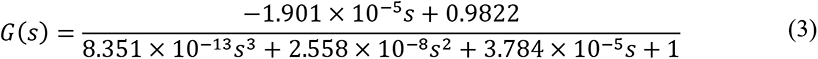

As shown in Figure 2c, the identified transfer function can model the system with a high accuracy. The closed-loop block diagram of the force sensor was shown in Figure 2a’. The force sensor stage displacement is detected and measured by the electrothermal sensor. The error signal was defined as the difference between the null position setpoint (i.e. 0 µm displacement) and the force sensor stage displacement measured by the electrothermal sensor. Using this error signal the controller generates a command signal which is then amplified by the voltage amplifier. The amplified command signal (*ν_a_*) and its inverse (–*ν_a_*) are separately added to the bias voltage and are applied to the opposite sides of the electrostatic actuator. This generates a nullifying force *F_n_* that pushes back the stage by *d_n_* in order to negate the induced external displacement *d_e_* caused by the external force.

The controller C(s) was designed with the MATLAB control system designer toolbox and is represented with the following transfer function:

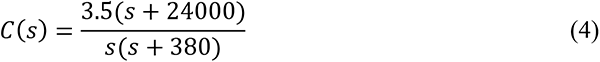

This controller has a large stability margin and accurate low-frequency tracking performance.

### Micro-positioner platform fabrication for in-vivo measurements

To maintain stabilization in the system and reduce the mechanical artifacts due to the breathing of the animal, a stabilizing platform was built (Figure 3). It consisted of two components, 1) A 3D printed base that was used to assemble the micropositioner system for the *In-vivo* measurements. This platform allowed to maintain the rat under inhaled anesthesia (isoflurane), with a warm pad and monitored vital signs during the experiments. The design and dimensions for the 3D printed base are detailed in Supplementary Figure 1. 2) A micro-positioner system to control the MEMS-CLFS. A high-precision linear stage with a 13 mm travel range (Newport, M-562-XYZ ULTRAlign) was used to mount a micrometric vernier (Newport, SM-13) that allowed movement control with a sensitivity of 1 µm, and with a differential micrometer with sub-micrometer resolution, and sensitivity of 0.07 µm (Newport, DM-13).

**Figure 3.**
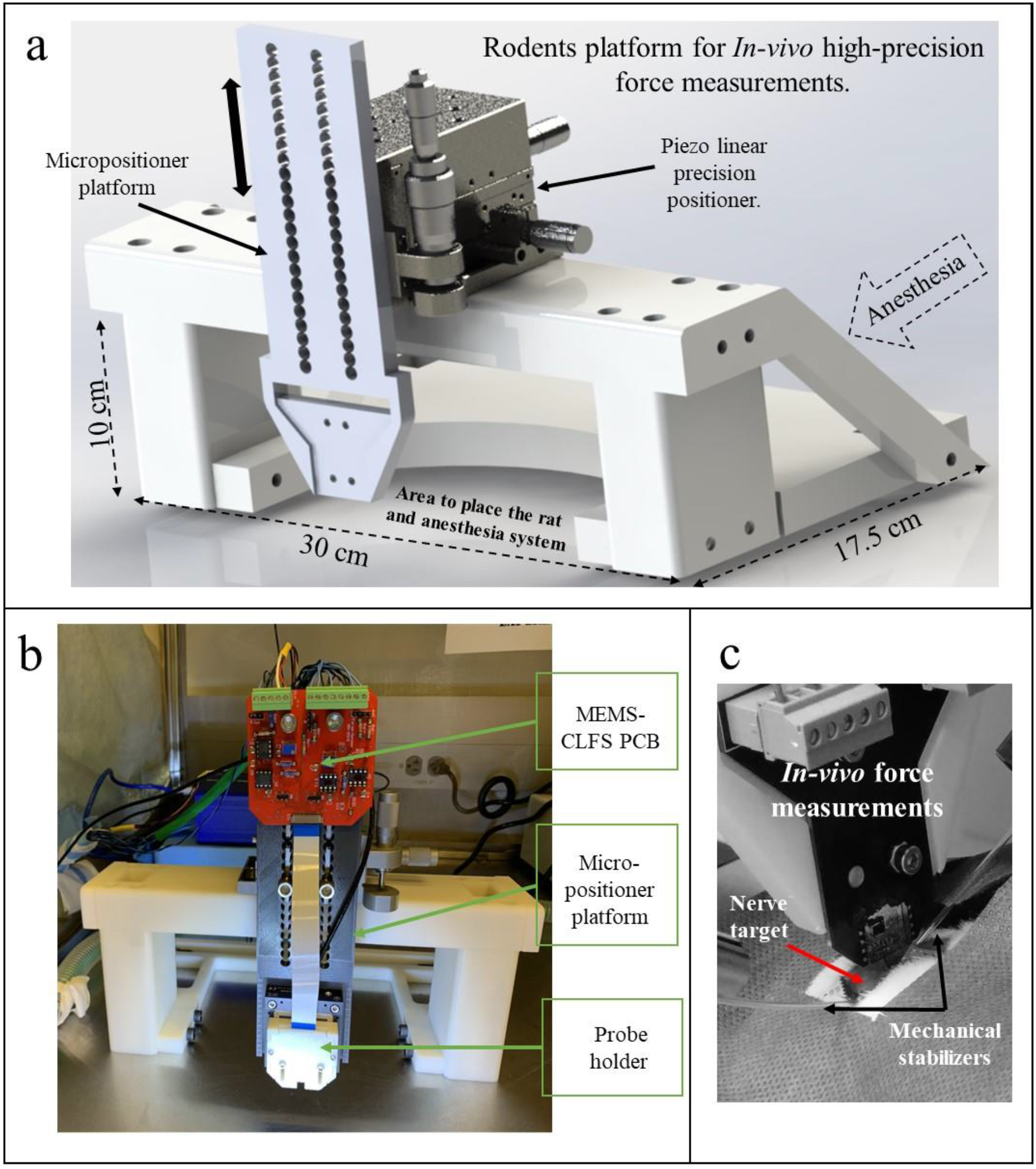
Design of rodent platform for *In-vivo* force measurements in peripheral nerves. a) Design of 3D printed platform for MEMS control and pneumatic micromanipulation. The double head black arrow indicates the z-displacements that allow the measurements: a dotted line arrow points to the area where the tubbing for anesthesia was assembled. b) Picture of plataform for measurements after the electronic components have been assembled, a mechanical micromanipulator allowed to get closer to the nerve target, while the automatized pneumatic system was used to maintain uniform velocities and depths. c) Picture of the MEMS probe when measuring from a nerve in the anesthetized rat, two glass stabilizers are pointed out, which allowed holding the nerve during the experiments. c’) image of the nerve with the tip of the probe while being inserted in the epineurium.

### Biomechanics of the epineurium

The procedures were performed in anesthetized rats, the sciatic or vagus nerve was exposed, and the animal was translated to the measurement area, which was conditioned with a warm pad, and physiological parameters monitor (Figure 4a). Mechanical stabilizers were used to maintain the nerve isolated from surrounding breathing artifacts. In the nerves were clearly identified under the microscopy those areas with high irrigation, so the measurements were obtained from i) more vascularized areas, where the tip was inserted between arteries (approx. 200µm far), and from ii) non-vascularized areas. The nerve inner composition of the vagus and sciatic nerve are schematized in Figure 4b. A representation of the MEMS-CLFS displacement during measurements is presented in Figure 4c.

**Figure 4.**
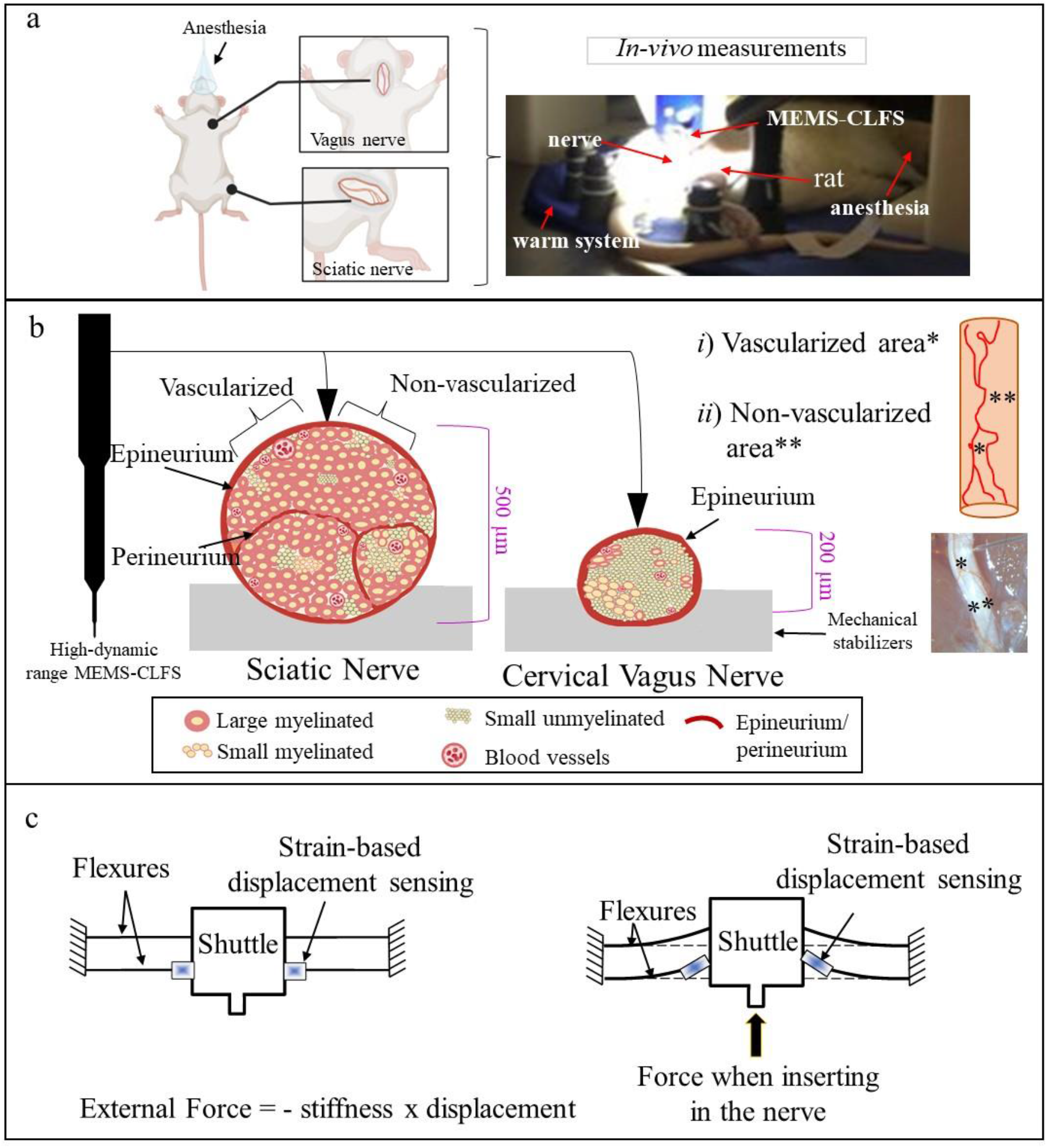
Experimental set-up for insertion force measurements. a) Animal semp. a cartoon and a picrure (left and right) show the location of the vagus and sciatic nerves (VN and ScN). The rat was under anesthesia while performing the measurements with the MEMS-CLFS held by a micromanipulation system. b) Schematic displaying the MEMS-CLFS and a representation of a ScN and VN cross-section. The modality and proportion of different nerve fibers (i.e. small and large myelinated, and non-myelinated) are different in these nerves. The measurements were performed in areas with dense vascularization and in areas where fewer blood vessels were visible (top-right, *i* and *ii*). c) Representation of mechanism of force measurements in the MEMS-CLFS by flexures displacement.

### Young Modulus of the Epineurium

The compressive stiffness or young modulus from the sciatic and vagus nerve were determined by the displacement of the force sensor with the speed of the rupture force of the epineurium, which was identified by a depression in the plot of displacement versus force. This measurement was possible to obtain due to the closed loop bi-directional mechanism of the MEMS (asterisks in Figure 5a). The measurements were obtained from the right and left nerves, and the results showed not statistical significance among the Young’s modulus of the epineurium of the right and left vagus and sciatic nerves, but there was among nerves (Figure 5 b-c, p<0.005). Tu further explore the intraneural elements that may explain these mechanical differences, we evaluated the density of collagen on these nerves. We obtained cross sections of the ScN and cVN and stained for collagen VI. Figure 5d shows confocal images where the higher density of collagen VI is ∼ 8.5-fold for the ScN in contrast with the VN. Furthermore, since the collagen was observed densely condensed around the blood vessels (perivascular collagen), the outer/inner diameter (OD/ID) ratios were calculated. The results showed a higher OD/ID ratio for the VN (∼2.3 vs 1.7 for the ScN), which indicates a higher condensation of collagen VI associated to the vasculature (graphics in Figure 5d).

**Figure 5.**
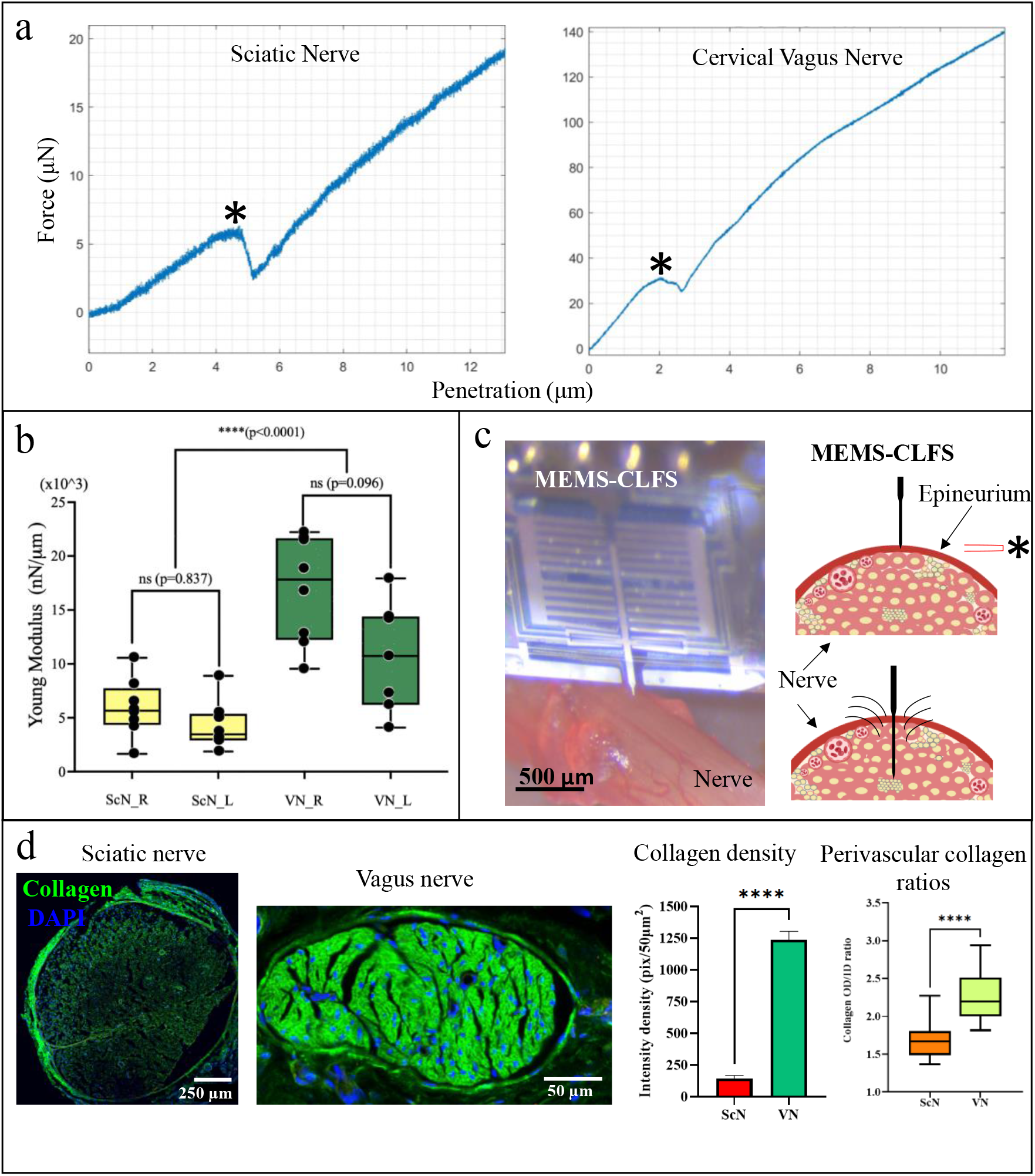
Effect of collagen in Young Modulus. High-dimensional MEMS force sensor allowed to elucidate the epineurium young modulus from the sciatic and cervical vagus nerve (SnN and VN, respectively). a) Two representative graphics of the insertion forces measured from the sciatic and cervical vagus nerve. The bi-dimensional capability of the MEMS allowed to evidence the rupture of the epineurium, which in turn allows to determine its young modulus (*), which is represented in (b) to compare right and left nerves (R and L). c) Picture and schematic representation of high-dimensional MEMS insertion and the rupture of the epineurium (*). d) Collagen VI staining (green) shows higher density for the vagus nerve in contrast with the sciatic nerve, with higher distribution around the blood vessels. The fluorescence was compared as intensity density per area (n=3 simples per nerve, 6 measurements each) and the relation of outer diameter/inner diameter (OD/ID) of the perivascular collagen was higher for the vagus nerve (graphic in the right). t student test (p˂0.005 for b and p˂0.0001 for d) was applied, n=7-9 measurements per nerve for a-c and n=3 measurements/animal, n=3 animals for d.

### Mechanical Properties of Extracellular Matrix of the Nerves

The sciatic and vagus nerves demonstrated different densities of collagen, which directly impact the mechanical properties of the extracellular matrix. Furthermore, different axonal composition, epineurium thickness, and irrigation are different for both nerves, which prompted us to evaluate matrix penetration forces. After the depletion of the probe due to the rupture of the epineurium, we measured the insertion force when penetrating the nerve. From the slope and linear equation, we calculate the force required to penetrate 25 µm of the nerve. The results showed statistically significant differences between the sciatic and vagus nerves, but not between the right and left nerves (Figure 6a-b, p<0.0001). The anatomical differences were evident when analyzing cross-sections from both nerves and contrasting with the marker vimentin (6c-d’), a structural intermediate filament, which density was also increased in the VN in contrast with the cVN.

**Figure 6.**
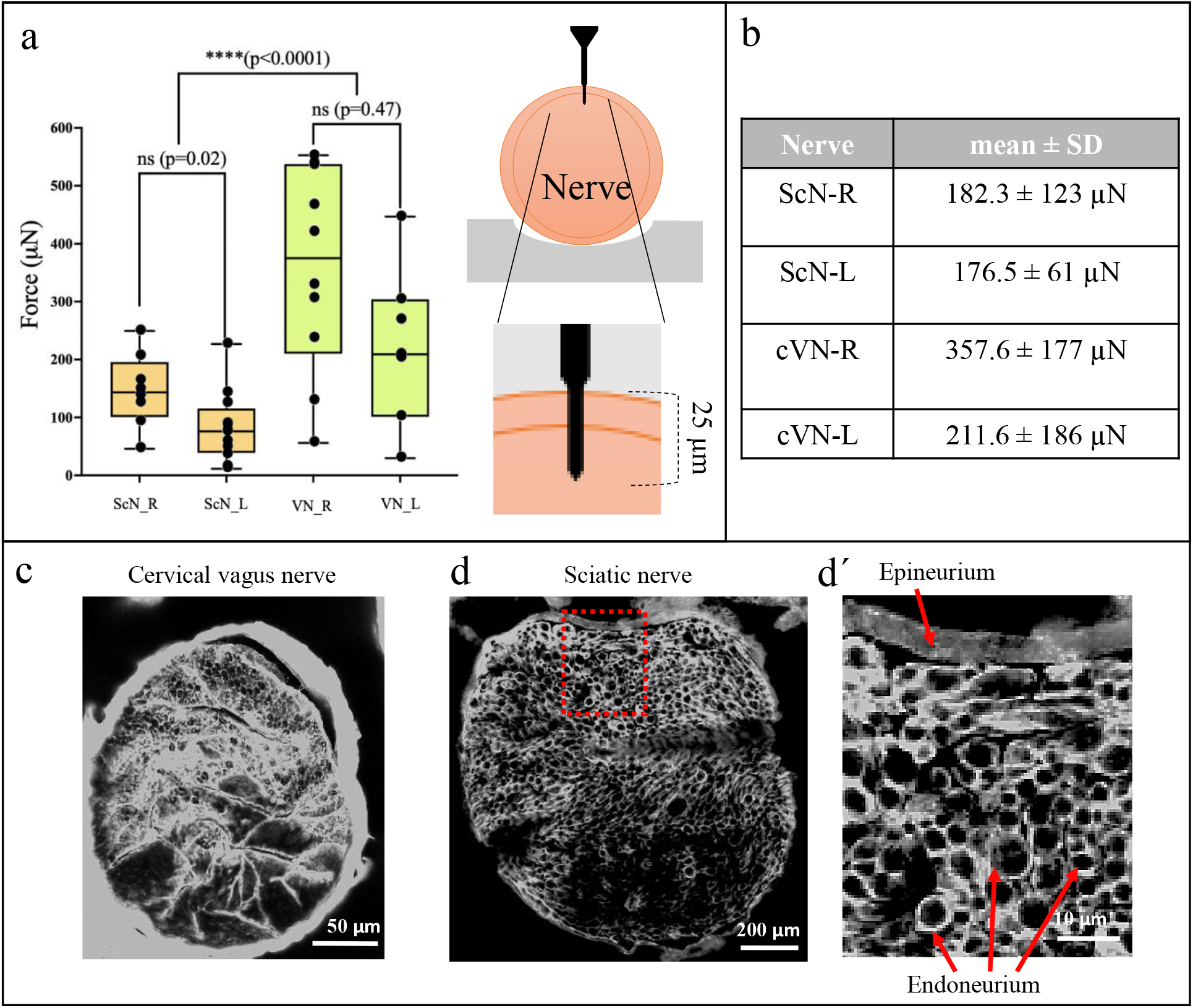
Insertion force required to implant a probe 25 µm through the epineurium. a) Graphics of all values (vascularized + less vascularized areas) of forces required to insert the probe 25 µm deep through the epineurium in the right and left sciatic and vagus nerves (ScN_R, ScN_L, VN_R and VN_L). A schematic representation is displayed on the right. b) mean ± SD of forces from a 95% of confidentiality criteria (p˂0.005) was applied, 8-12 measurements per nerve. c-d) confocal images of cross-sections obtained from the cervical vagus nerve and one fascicle of the sciatic nerve. Epineurium and endoneurium are stained with vimentin, d displays the square outlined in d.

### Biomechanics of the Blood-Nerve Barrier

The nerves are highly vascularized, blood vessels are distributed internally and run in the surface of the epineurium and in the inner layers of the perineurium and the endoneurium. Our studies of collagen distribution demonstrated higher density around blood vessels, which prompted us to further explore the BNB density in both nerves. We evaluated the stiffness of the epineurium near the vascularized areas (< 200µm far) and in non-vascularized areas (> 200µm far). The results showed that right and left sciatic and vagus nerves have statistically significant differences in stiffness when comparing vascularized versus non-vascularized areas (Figure 7a). We then analyzed CD31 expression, a marker for the endothelial cells associated to the BNB and measured the vasculature per area. The vasculature forming the BNB was associated to collagen (perivascular collagen), we refer these structures CD31+ blood vessels-collagen as BNB clusters, due to their clusterized appearance with low magnification microscopy. Our analysis demonstrated that the VN has ∼8.6 fold density of BNB clusters in comparation with the ScN (Figure 7b). Furthermore, the CD31+ expression in larger vasculature presented different patterns for the VN and ScN, with fenestrated-like (discontinuous) epithelium in the ScN, and continuous in the VN (Figure 7c).

**Figure 7.**
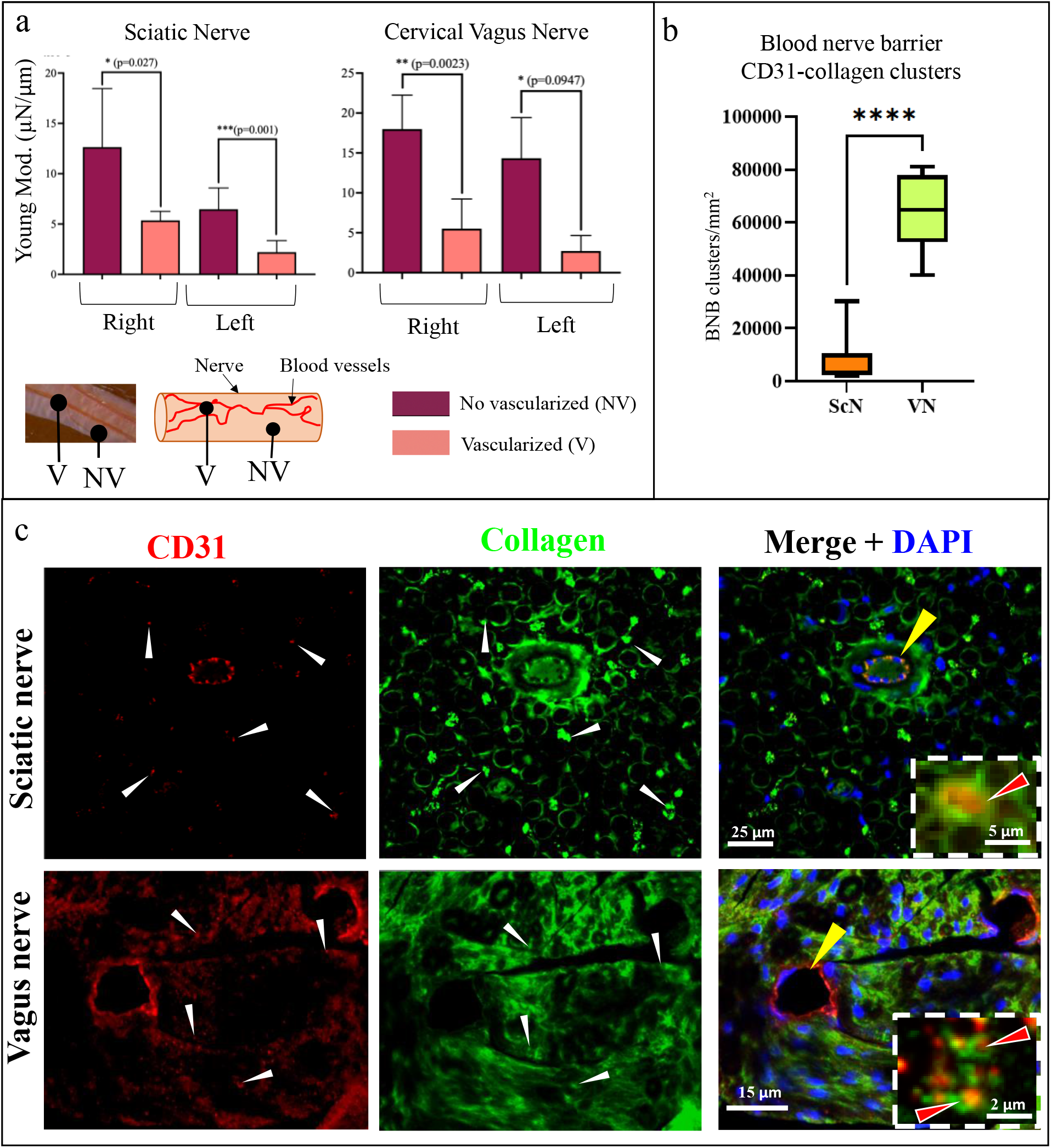
Effect of blood nerve barrier in Young modulus of peripheral nerves. a) Stiffness of the right and left sciatic and cervical vagus nerves. No statistical diference was observed between the right and left groups for NV sciatic nerve (p=0.20), V sciatic nerve (p=0.16), NV vagus nerve (p=0.33) nor V vagus nerve (p=0.42). A nerve picture and a schematic representation is displayed in the bottom. b) Blood nerve barrier clusters were defined as blood vessels CD31+ surrounded by collagen, their distribution per area was defined. The histological immunostainings are presented in (c), BNB clusters are pointed with whita arrowheads. Large blood vessels with different patterns of BNB were observed in both nerves (yellow arrowhead). The squares with dotted lines show examples of BNB clusters (Orange arrowhead). One way ANOVA p˂0.005, n=15-17 values per nerve.

### Predicting penetration force for nerve interfacing

The penetration force measurements, displacement and area of the probe lead us to explore more mechanical properties of these nerves by applying physics principles. The force graphics in Fig. 5a clearly shows the epineurium puncture and young modulus, these values were used to calculate the stress and strain of the nerves at different phases, including ultimately, yield and failure (Figure 8 a-b). The ratios of these values were different among nerves but showed a direct strain/stress correlation of 0.4 (Table 1), which was taken into account to estimate a constant factor, and the root square error (RSE=0.2, Figure 8c-d) validated this predicted correlation with the original data from the nerves.

**Figure 8.**
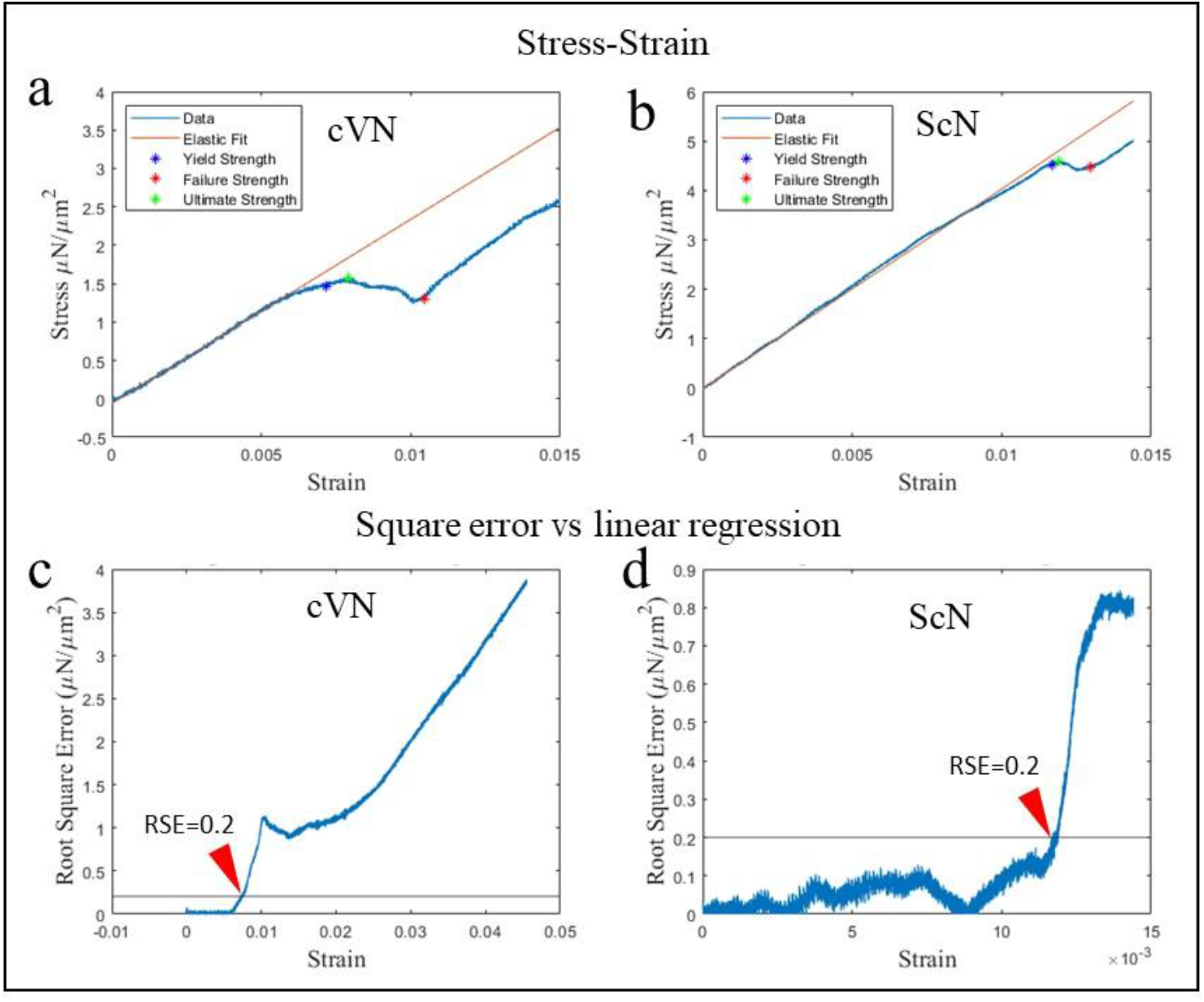
Elastic-Plastic behavior of the epineurium shows distinct yield, ultimate, and failure points. A comparison of elastic model versus *in vivo* mechanical response for representative stress-strain curves for cVN and ScN. Panels a and b compare the elasfic and models and indicate where plastic deformation begins, the ultimate strength of the epineurium, and the puncture point on the curve. Panels C and D show the Root Square Error (μN/μm^2^) between the linear elastic fit and the actual data. An RSE of 0.2 indicated significant deviance from the elastic fit and therefore the beginning of the plastic deformation region.

**Table I.**

|  | Ratio (cVN/ScN) | Ratio of diameters (cVN/ScN) |
| --- | --- | --- |
| STRESS | Ultimate Stress Ratio | 0.3424 |
|  | Yield Stress Ratio | 0.3914 |
|  | Failure Stress Ratio | 0.2882 |
| STRAIN | Ultimate Strain Ratio | 0.5399 |
|  | Yield Strain Ratio | 0.6206 |
|  | Failure Strain Ratio | 0.6477 |
|  | Diameter Ratio | 0.4 |

The constant correlations among nerves were used to define an equation that integrates the stress, ultimate stress, the nerve diameter and the area of the penetration probe to predict the required force to penetrate a nerve. The equations (5-7) allowed us to define (8), which was validated with real values obtained from the sciatic and vagus nerves published by others, and proved to be accurate^22, 23^.

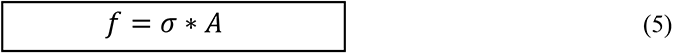

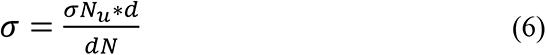

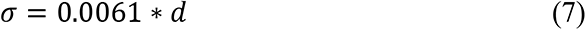

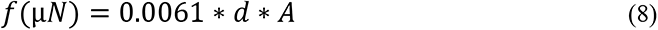

*f =* force required for epineurium puncture

σ= stress for epineurium penetration

*A*= probe area

*σNu =* nerve ultimate stress

*dN* = nerve diameter

## Materials and Methods

### Animal use

#### Ethics statement

All the protocols and surgical procedures were designed to prevent animal discomfort and suffering. These were approved by The University of Texas at Dallas, Institutional Animal Care and Use Committee (IACUC, protocol No.14-09), and follow the guidelines provided by the National Institute of Health (NIH).

### Surgical procedures

A total of seven male Sprague Dawley rats (300-350 g; Charles River, Wilmington, MA) were used for the experiments. The animals were anesthetized with vaporized isoflurane (2%) in a constant oxygen flux (2 L/min) delivered by a calibrated vaporizer and maintained throughout the experiment. The animal temperature was maintained with an electrical warm pad. Anesthesia and vital signs were monitored constantly with standard methods throughout the experiment. Individual right and left nerves studies included: the sciatic nerve (ScN) and the cervical vagus nerve (cVN). The *Sciatic nerve (ScN)* was exposed by a 4 cm longitudinal incision on the animal’s hind limb (right and left) from hip to knee following the femur. The biceps femoris and vastus lateralis muscles were separated to visualize the nerve. The ScN was identified as a ̴ 500µm diameter fasciculate nerve lying longitudinally between these muscles. It was cleaned from connective tissue and carefully placed on a stabilizing platform in order to maintain mechanical stability during the measurements. The *cervical vagus nerve (cVN)* was exposed by making a longitudinal medial incision in the anterior part of the neck at the cervical level, using the anterior sternum as an anatomical reference. The sternomastoid muscle was separated in oblique orientation to the midline and the cVN was identified lateral to the carotid artery. Surgical retractors were used to maintain the separation of the muscles. During the nerves exposure, efforts were taken not to disrupt the epineurium. The nerves and surrounding tissue were kept hydrated with sterile 1x phosphate buffered saline (PBS, pH 7.2) for the duration of the experiment.

### MEMS-CLFS microfabrication

The force sensor used in tissue indentation tests was fabricated using the silicon-on-isolator multi-user MEMS process (SOI-MUMPs) offered by MEMSCAP Inc. The three-mask process utilized a silicon-on-isolator wafer comprising a 25-μm-thick Si device layer, a 2-μm-thick buried-oxide, and a 400-μm-thick handle layer.

The process started with diffusion doping of the device layer to enhance electrical conductivity. Next, the electrode layer forming the pads and routing for the electrical connections was deposited and lifted off. After that, the device layer was patterned and dry-etched to form the force sensor body. Finally, the Si handle layer and box layers were etched using dry and wet etching techniques, and the devices are released. Figure 1 shows an SEM micrograph of the microfabricated force sensor. The microfabricated devices were then glued and wire-bonded to a custom-design printed circuit board (PCB) for characterization. The force sensors were inspected to ensure the quality of fabrication using scanning electron microscopy.

### In-vivo measurements

All the measurements were performed inside a Faraday cage covered with a fabric cloth to prevent electrical or mechanical interference from the environment. Force measurements were obtained from the right and left cVN and ScN. Three to five measurements were obtained per animal. The rat was placed under the micropositioner stage, the nerve to study was oriented perpendicular to the positioner and maintained stable by using micromanipulator arms. An M320 Leica surgical stereoscope was used to guide the measurements and to obtain images and videos. The force sensor’s PCB was put in a micropositioner platform, which consisted of a custom-made 3D printed casing that was assembled on a PI P-622.1CD Piezo Linear Precision positioner. The positioner was fixed on a custom-made 3D printed holder that could be moved by a Newport M-562-XYZ Linear Stage in X, Y, and Z directions.

### Immunohistochemistry

At the end of the studies, the animals were euthanized with an overdose of sodium pentobarbital (120 mg/kg, intraperitoneal). The ScN and the cVN were harvested and fixed in cold 4% paraformaldehyde diluted in phosphate-buffered saline (PBS, pH 7.2) for 24 h, cryoprotected in gradients of 10, 20 and 30% sucrose in PBS, embedded in optimal cutting temperature (OCT) media, and cut in 35 μm cross-sections in a cryostat. The sections were rinsed, blocked, and incubated with primary antibodies as described previously ^24^. Antibodies from Abcam^®^ were used, against CD31 (rabbit, ab222783) and monoclonal Alexa Fluor^®^ 488-collagen VI (ab200429, and the secondary antibodies coupled to Alexa Fluor 594^®^ (goat anti-rabbit, ab150080). The slides were mounted with Fluoroshield-DAPI (ab104139). The sections were imaged in a confocal microscope (Nikon, eclipse Ti^®^). Collagen fluorescence intensity was calculated per area and measured blood vessel ratios as outer diameter/ inner diameter (OD/ID) ratios. The blood vessels CD31+ were measured by area for statistical comparisons.

### Statistical Analysis

The force and elastic modulus differences between groups were evaluated using one-way ANOVA. Unpaired student t-test was used for global value comparison and histological analysis. The comparisons were considered statistically significant as p˂0.005 or p˂0.001, indicated for individual analysis. GraphPad Prism software version 9.1.2 and MATLAB R2020a were used to prepare graphics and perform statistics.

## Discussion

The engineered design of intraneural interphases for naturalistic sensory-motor prosthesis requires high precision and special resolution while minimizing implant trauma^25, 26^. The peripheral nerves have been treated as homogeneous structures when designing neural interfaces, and the anatomy and physiology demonstrate that is required of microscale customization on the design to minimize implant damage that compromises the function^27^. Here we report the use of a high dynamic range closed loop MEMS to define the biomechanics of two nerves with different modalities, somatic and autonomic (ScN and VN, respectively). The membrane structure of the nerves and axonal composition directly affect the biophysics of peripheral circuits, and differ in their mechanical and conductive properties due to different proportions of collagen and lipids (i.e. extracellular matrix, epineurium thickness and axon myelinization)^28–31^. For instance, myelinated fibers have higher conduction velocities than unmyelinated, and the proportion of these fibers vary depending on the nature of the nerves^32, 33^. Somatic nerves carry sensory and motor fibers associated with proprioception, pressure, pain and temperature with more myelinated axons and with nerve conduction velocities ranging from 10-120 m/s, these include Aα, Aβ, Aγ and Aδ fibers. In contrast, autonomic nerves carry mainly unmyelinated fibers with lower conduction velocities ranging from 0.3 to 3 m/s ^28, 29, 34^.

The recently growing field in bioelectronic medicine demands advances in neural interface methods to miniaturize device sizes and engineer soft materials to reduce tissue trauma and optimize function^35, 36^, but less attention receives the nerve composition individuality. In this work we focused on exploring the mechanical properties of different nerve modalities, somatic with a higher number of myelinated fibers and autonomic with primarily unmyelinated fibers. The sciatic nerve is a model used to study somatic neves^24, 25^, while the vagus nerve has gained attention as a main autonomic nerve connecting the brain with peripheral organs and is a main focus for neuromodulation as a treatment for bioelectronic medicine^11, 13, 37, 38^.

The measurement of insertion forces to penetrate the sciatic nerve has been reported previously in rabbits as ∼7.2-71.8 mN for a range of 50-200μm diameter sharp needles ^39^, while in our previous studies *in-vivo*, in the rat sciatic nerve we tested different velocities and inclinations in the penetrating shank to define a range of penetration forces of 20-120 mN for a shank 2 mm of length, 30 μm thickness and 100 μm width ^25^. Our previous data on determining the force required to implant intraneuronal electrodes *in-vivo*, in the rat sciatic nerve allowed us to determine values between 10-60 mN and 80-125 mN for angled and straight insertion (with respect to the nerve surface). However, the measurements were limited to mN resolution and did not take into account the irrigation of the measurement area^25^. Other studies promised to define the epineurial and perineurial biomechanics of the rabbit sciatic nerve ^40^. They report an insertion force of 3.3 ± 0.5 and 40.0 ± 8.2 mN and Young’s modulus of 3.0 0.3 and 0.4 0.1 MPa for the epineurium and perineurium, respectively, however the technology is an open loop commercial load transducer with resolution in the order of mN, not enough to define intracellular matrix properties. Their studies provide evidence of the effect of sharp and flat tip insertion; however their studies were done in explanted nerves, and in a time window of twenty-four hours. In our experience, after five minutes of explant, the biomechanics in the nerves change, which prompted us to perform our studies *in vivo*.

With the MEMS-CLFS used here, we further expanded our previous studies, achieving higher resolution on the range of µN and compared between irrigated and less irrigated areas in the nerves. The penetration tip probe used had 2x10 µm of surface area and 11 µm in length, in contrast with others of 30x100 µm and up to 300µm of length^41^, which allowed high precision control over the nerve target, and the closed loop dynamic control allowed to compensate micromotion while measuring. The first approach was the epineurium contact, which was represented as an elastic material. The epineurium with clear elastic behavior allowed to determine yield, failure and ultimate strain of the sciatic as 0.012, 0.03, 0.012 µN/µm^2^, and the vagus nerve 0.007, 0.010, 0.008 µN/µm^2^, respectively, while the yield, failure and ultimate stress was determined for the sciatic nerve as 4.515, 4.480, 4.60 µN/µm^2^, and for the vagus nerve 1.456, 1.307, 1.574 µN/µm^2^ respectively (Figure 8). This mechanical behavior allowed to determine the young modulus of both, sciatic and vagus nerve, which presented statistically significant differences (p˂0.005, Figure 5a-b). This value is remarkedly critical for the design of electrodes, as represents the force required to break into the epineurium, and our data highlights the necessity of customize these forces for different nerves. The graphics of force-displacement allowed to identify the force required to penetrate the epineurium as a flexure (Figure 5a). This measurement was possible due to the bi-directional closed loop of the MEMS probe, which allowed to compensate micromotion movements and adapt to the dynamic environment. Previously the use of MEMS-based force sensors complemented with atomic force microscopy (AFM)-based nanoindentation allowed to measure nanoscale properties from individual collagen fibers ^41–44^ with high resolution in the range of nN, however such technology has been suitable for *in-vitro* studies, and it is well known that explanted tissue modifies its mechanical properties. To our knowledge, nerves and other neuronal structures mechanical properties have been defined from explanted tissue and no detailed information is available to take in account the epineurium breakage and fibers composition dependence^8, 10, 40, 45^. We provided these values measured directly from the alive animal and demonstrated mechanical differences between somatic and autonomic nerves.

While the mechanical properties of the epineurium were determined, the internal fiber composition of the nerves demand differential measurements. After the MEMS-CLFS penetrated the epineurium, the probe was advanced to measure intra-fiber biomechanics. The data showed that the internal biomechanics of the ScN and VN were significantly different, which reflects the fiber composition with predominantly large myelinated fibers for the ScN and small unmyelinated for the VN (from 176-182 µN for sciatic and 211-357 µN for the vagus nerve, p˂0.005). The micrometric surface area of the probe and its high sensitivity provided the opportunity of differentiating between highly vascularized areas and less irrigated nerve surfaces. The criteria was measuring less or more than 200 µm far from blood vessels, for vascularized and no vascularized areas, respectively. The results showed that for both, the sciatic and the vagus nerve the penetration forces vary between vascularization degrees (Figure 7a). The difference in mechanical properties between the ScN and VN lead us to investigate the extracellular matrix in more detail. We stained collagen VI, a well-known highly abundant structural protein in the nerves, and CD31, a marker expressed in endothelial cells associated with the blood nerve barrier. We demonstrated that the VN had a ∼ 8.5-8.6 fold density of collagen VI and vasculature (CD31+) per area than the ScN. Furthermore, the collagen VI formed dense clusters around the BNB, with OD/ID ratios of 2.3±0.3 and 1.7±0.1 for the VN and ScN, respectively. Collagen is a structural component ^30, 41^, and its distribution suggests a relation to the BNB, which with higher density in the VN explains its higher penetration forces and different Young’s modulus.

The BNB is considered the second most selective vascular system after the blood-brain barrier, and here we demonstrate that has different properties in autonomic and somatic nerves. In general, it is known that structurally consist of an outer collagenous epineurium, inner perineurium myofibroblasts defining the endoneurium, which envelops myelinated and unmyelinated axons embedded in a looser mesh of collagen fibers^16, 46^. The microstructure of the nerves has medical implications, since its integrity is crucial for nerve health and if disrupted leads to neuropathology (i.e. Guillen Barre syndrome)^15^. Different degrees of irrigation, involve blood vessels in the surface and inner environment, directly impacting on the mechanical properties ^46, 47^. While different nerves share similar characteristics, the differences in the epineurium thickness, proportion of axons and diameters, myelinization, and the degree of irrigation, provide them with different mechanical properties ^25, 27, 48^.

Our data showed that biomechanical properties of somatic and autonomic nerves differ due to three main elements: i) epineurium Young’s modulus, ii) collagen density, and iii) BNB. By having mechanical values differential for nerves, we reasoned that an equation may help to determine the force required to penetrate a nerve just by knowing its elastic modulus measured by no invasive methods. Here we propose an equation that will allow this calculus by incorporating a factor value, the area of a probe, and the Young’s modulus of a nerve.

Overall, our MEMS-CLFS system allowed us to evidence the differences in the mechanical properties of the ScN and VN, which could be explained by the differences in collagen and BNB density and distribution. We propose a mathematical algorithm to deferentially predict penetration forces of somatic and autonomic nerves based on the stress, ultimate stress, the nerve diameter and the area of the penetration probe to predict the insertion force.

In conclusion, we used a high precision MEMS-CLFS technology to demonstrate *in-vivo* that somatic and autonomic nerves differ in their BNB biomechanics. These differences should be considered in the development of precision technology of penetrating interfaces. Our MEMS-CLFS was critical to define with high resolution the biomechanics of the ScN and VN, and its fundamentals on the principle of force balancing using an electrothermal closed-loop control in a wide range of resolution, which is critical when dealing with the mechanical characterization of samples with wide ranges of stiffness.

## Aknowledgements

This work was funded by NIH NINDS 1R01NS124222 -01.

MAGG and MR acknowledges the technical support of Danny Lam and Atefeh Ghazavi at the University of Texas at Dallas.

## Author contributions

Experimental design: MAGG, MBC, MM

*In-vivo* studies: MAGG, HA, MBC, MM

Mathematical model: DL

Analyzed the results: MAGG, HA, MRO, SORM

Reviewed the manuscript: MAGG, HA, MM, MBC, DL, SORM, MRO

Wrote the paper: MAGG, HA

## Conflict of Interest

The authors declare no conflict of interests.

## Notes

### Competing Interest Statement

The authors have declared no competing interest.

